# Substrate stiffness controls the cell cycle of human mesenchymal stem cells via cellular traction

**DOI:** 10.1101/2021.11.19.469207

**Authors:** Sanjay Kumar Kureel, Shatarupa Sinha, Purboja Purkayastha, Sarah Barretto, Abhijit Majumder

## Abstract

The microenvironment of human mesenchymal stem cells (hMSCs) regulates their self-renewal and differentiation properties. Previously it was shown that hMSCs remained quiescent on soft (0.25 kPa) polyacrylamide (PA) gels but re-entered into cell cycle on a stiff (7.5 kPa) gel. However, how cells behave on intermediate stiffness and what intracellular factors transmit mechanical changes to cell interior thereby regulating cell cycle remained unknown. In this work we demonstrated that PA gels between 1 and 5 kPa act as a mechanical switch in regulating cell cycle of hMSCs. By experiments on cell-cycle exit and re-entry, we found that hMSCs demonstrated a sharp transition from quiescence to proliferation between 1 and 5 kPa. Further studies with ROCK inhibitor Y-27632 revealed that contractile proteins, but not cell spread area, accounts for the sensitivity of hMSCs towards substrate stiffness and hence correlates with their changes in cell cycle. These observations therefore suggest that substrate stiffness regulates hMSC proliferation through contractile forces as generated by cellular contractile proteins in a unique pattern which is distinct from other cell types as studied.

**Graphical abstract:** 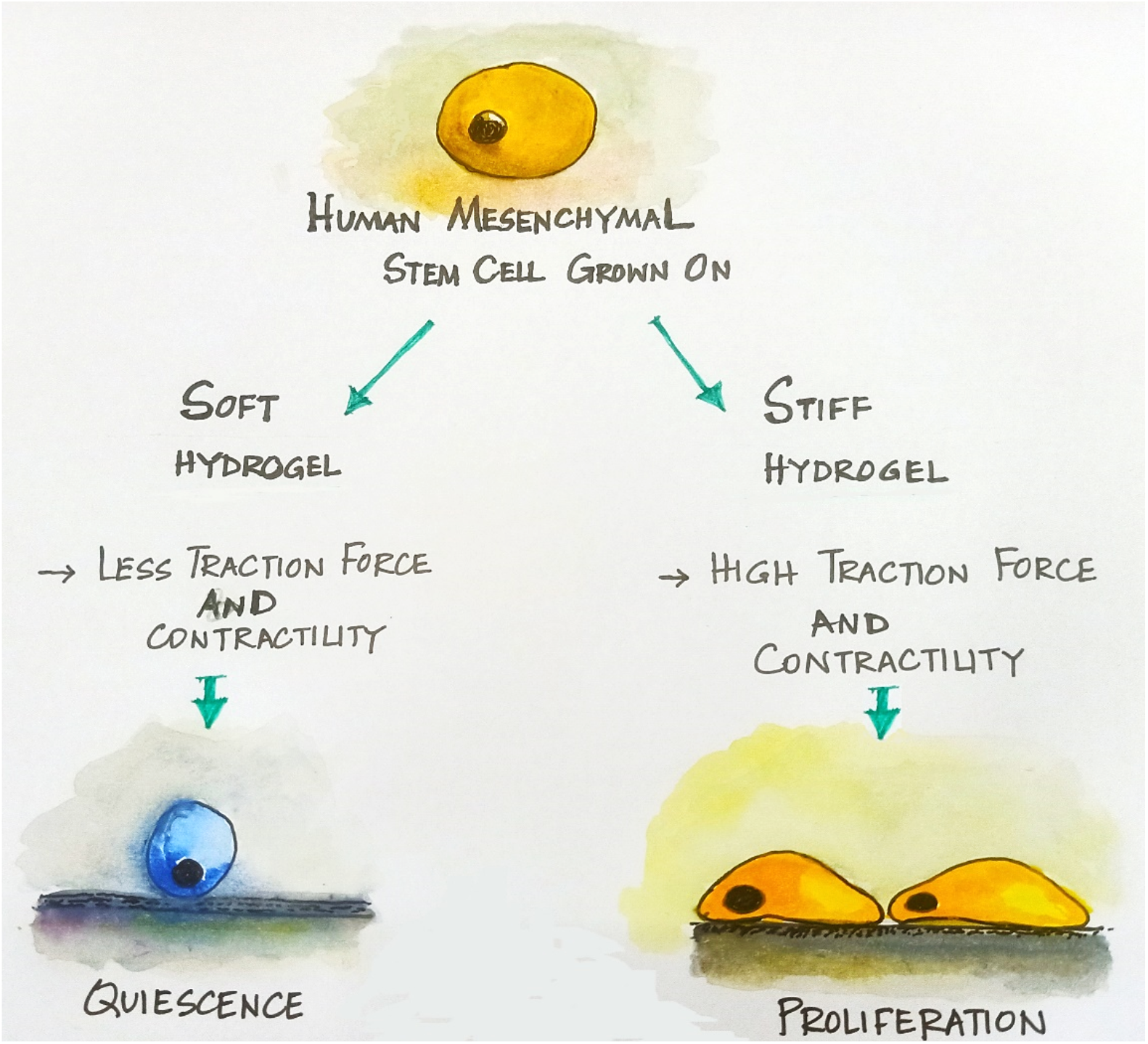

## 1. INTRODUCTION

Rigidity of the extracellular matrix (ECM), along with other chemical and mechanical cues, controls proliferation of the adherent cells. Such mechanobiological control is crucial during development and homeostasis[1]. Loss or malfunctioning of such control leads to various pathological conditions[2][3].

To understand the role of ECM rigidity on proliferation, *in-vitro* studies have been conducted on polyacrylamide (PA) hydrogels whose elasticity can be tuned easily. Winer et al. (2008) showed that human Mesenchymal Stem Cells (hMSCs) on soft gels (0.25 kPa PA) became round, exhibited reduced DNA synthesis, and attained reversible cell cycle arrest called quiescence [4]. However, these quiescent hMSCs could proliferate or differentiate when transferred on stiff gel (7.5 kPa) in presence of appropriate media. Interestingly, some other cell types such as NIH-3T3 fibroblasts and aortic endothelial cells proliferate even when grown on gels (< 0.25 kPa) soft enough to induce round morphology[5]. The questions remain (a) how do hMSCs behave within the intermediate stiffness of 0.25 kPa and 7.5 kPa?, (b) why different cell types behave differently?, (c) does cell cycle entry and exit occur around the same stiffness value or is there a hysteresis?, and (d) what mediates changes in substrate stiffness to regulate cell cycle?

We have addressed these questions in this paper using three cell lines – bone-marrow derived hMSCs, immortalised mouse myoblasts (C2C12) and mouse fibroblast cells (3T3) cells. Each cell line showed different response towards the range of substrate stiffness (20 kPa to 0.5 kPa) investigated in this study. While hMSCs displayed a sharp transition from proliferating to quiescence state between 5 and 1 kPa gels, C2C12 displayed a gradual decrease and 3T3 cells remained insensitive to substrate rigidity. Our investigation further revealed that cellular contractility, but not cell spread area, accounts for the cellular sensitivity towards substrate stiffness. We confirm that contractile proteins act as mechanosensors, translating changes in surrounding matrix to cell interior, thereby regulating cell cycle events in hMSCs.

## 2. MATERIALS AND METHODS

### 2.(i) Substrate preparation

This protocol is adapted from Tse and Engler [6]. Briefly, by mixing 40% polyacrylamide and 2% bis-acrylamide solution[7], polymerized gels were conjugated with sulfo-SANPAH (G-Bioscience) in HEPES buffer (pH = 8.5) using UV cross-linker (Genetix, 312 nm) followed by collagen (50 μg/ml) (Invitrogen, A1048301) coating[8].

### 2.(ii) Cell culture

Bone marrow-derived human MSCs were (Lonza,Cat #PT-2501, Lot #482966) maintained in low-glucose DMEM medium[8]. C2C12 and NIH-3T3 cells were maintained in their respective high-glucose DMEM growth media[9][10].

### 2.(iii) Fold Change

Images at final time point (nearing confluency) were acquired after Hoechst staining. Cells were counted in each frame (ImageJ) and the average was multiplied by total area of gel and divided by area of single frame to get the total cell count in gel. The fold change of proliferation was calculated as (*N*_*t*_ − *N*_0_)/*N*_0_, where *N*_*t*_ is the cell count at a particular time ‘*t*’, and *N*_0_ is the count at the initial time (i.e., after 4 hrs of seeding). Note that negative values of fold change indicate growth arrest or quiescence, and positive values signify growth.

### 2.(iv) Cells synchronization at G0, quiescent phase

This protocol is adapted from Arora et al.[11]. Briefly, to warm HG-DMEM (250 ml), methyl cellulose (10 gm) was added while stirring at 800 rpm. This was diluted with HG-DMEM (250 ml) and kept overnight at 4°C on stirrer ∼800 rpm. After centrifugation at 4°C, trypsinised cells were added to the supernatant, kept for 48 hrs and centrifuged at 1400 rpm to obtain a clear cell suspension.

### 2.(v) Cell-cycle re-entry

Followed by synchronisation (G0 phase), cells were collected and seeded on PA gels of varying stiffness. Images were captured to determine fold change values.

### 2.(vi) BrdU assay

48 hrs post-seeding, cells were incubated with BrdU reagent (Invitrogen, 1:100) for 4 hrs at 37°C with 5% CO_2_. This was followed by fixing (4% PFA), permeabilizing (0.5% Triton-X), denaturing (2M HCl), blocking (1.5% BSA), and incubating with primary antibody (Invitrogen, B35128, 1:100). Images were captured by EVOS-FL Auto and analysed with ImageJ.

### 2.(vii) Traction force microscopy

A hydrophobic coverslip with 20 μl gel solution containing red fluorescent beads (50:1, diameter 1μm, Fluka) was buried underneath a pre-solidified gel, both having same stiffness values. Phase contrast (for cell boundary) and RFP images (for stressed condition) were captured by EVOS FL Auto at 24 hrs post-cell seeding. Finally after adding 2% Triton-X to cells, RFP images were captured (unstressed condition). All three images were used to calculate traction force by MATLAB code[12].

### 2.(viii) Calculation of cell area

Images of hMSCs were captured using EVOS FL Auto microscope (Life technologies, USA) at 10X and analysed for their spread area using ImageJ (National Institute of Health, Bethesda, USA) software.

### 2.(ix) Inhibitor studies

ROCK inhibitor Y-27632 was used at varying doses – 1, 2, 5, 10 and 20μM on hMSCs grown on 5 kPa gel and incubated for 96 hrs (fold change) and 48 hrs (TFM). MEK-inhibitor PD98059 was used at varying doses – 1, 5, 10, 25 and 50μM on hMSCs grown on 5, 10, 20 kPa gels and TCP for 48 hrs.

### 2.(x) Immunofluorescence and confocal microscopy

Cells were fixed (4% PFA, 10 mins) permeabilized (0.5% Triton X-100, 15 mins) and blocked (1.5% BSA, 30 mins). They were incubated in phospho-ERK antibody (Cell signaling #9101) overnight followed by Alexa Fluor 488 conjugated secondary antibody (1:1000) and Hoechst 33258 nuclear stain (1:10000) for 2 hours. Images were captured using LSM 780 confocal microscope (Zeiss) at 63× magnification. Integrated mean density of the phospho-ERK signals were quantified by Fiji ImageJ software.

### 2.(xi) Statistical analysis

All data were expressed as average ± standard error (SE). Two-tailed and unpaired Student t-test was used for p-values.

## 3. RESULTS

### 3.(i) Cell cycle exit by hMSCs between 1 and 5 kPa PA gel

#### (a) Proliferation assay

Fold Change in cell number showed the biphasic response of hMSCs to substrate stiffness r (Fig.1b & *Supplementary Fig.1a*). They were (a) quiescent on 0.5 and 1 kPa gels, and (b) proliferating on gels of higher rigidities (5 kPa and higher) (Fig. 1a&b). On 5 kPa gel, hMSCs proliferated twice as much compared to 1 kPa (Fig. 1b). 2 kPa was the intermediate stiffness between growth-arrest and proliferation. There was no difference in cell proliferation on 5 and 20 kPa, signifying saturation of the rigidity effect.

**Figure 1.**
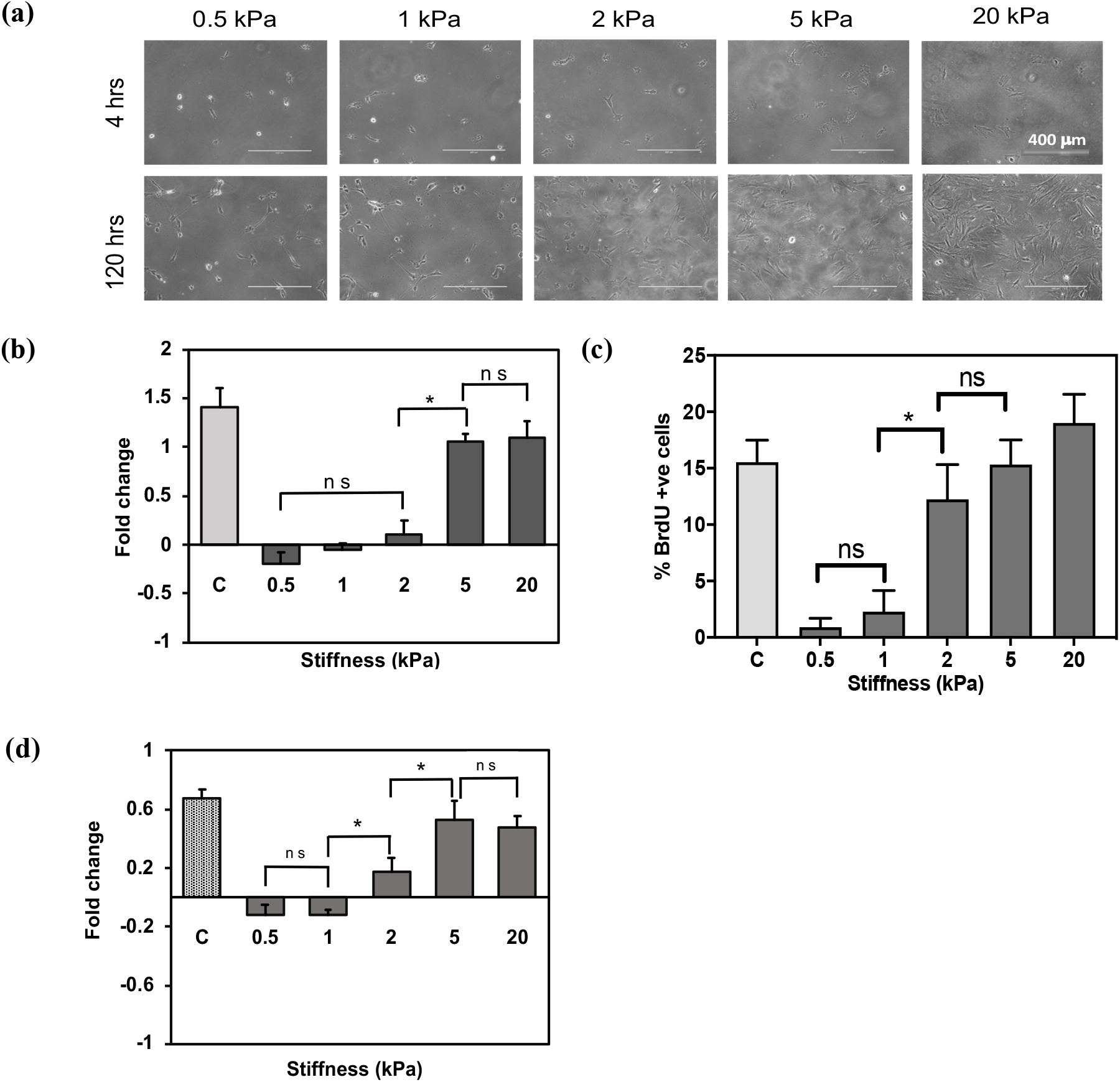
Effect of substrate stiffness on cell cycle exit & re-entry of hMSCs. (a) Representative images of hMSCs on substrates with various stiffness at different time points. Scale bar is 400 μm. (b) Fold change vs stiffness plot at 72 hrs. The experimental conditions are compared with the TCP (denoted as C) as control. (c) Plot of percentage of BrdU positive cells vs stiffness and control. (d) Fold change vs stiffness plot for cell cycle re-entry of hMSCs in 72 hrs. Results are represented as mean + SE, where n = 300 cells, N = 3 independent experiments, * p < 0.05 (Student’s unpaired t-test) and ns is non-significant.

#### (b) BrdU assay

To further confirm that lower fold change values obtained on soft gels were indeed due to reduced cell proliferation, BrdU assay was performed. Results showed that on 2, 5 and 20 kPa, 12 – 20% of cells were BrdU positive i.e. in ‘S’ phase (proliferating), whereas on ≤ 1 kPa gels less than 3% of cells stained positive (Fig.1c & *Supplementary Fig*.*2*).

### 3.(ii) Cell cycle re-entry by hMSCs between 1 and 5 kPa PA gel

We next addressed whether stiffness dependent cell cycle exit and re-entry has any hysteresis. hMSCs were found to re-enter into cell cycle at ∼2 kPa or higher (Fig. 1d & *Supplementary Fig. 3*) and continued to remain quiescent on ≤ 1 kPa. Taken together, these results indicate that both cell cycle exit and re-entry from quiescence phase are controlled by ECM stiffness, and there is no significant hysteresis.

### 3.(iii) Proliferation of C2C12 increases with substrate stiffness, 3T3 remains unresponsive

To investigate the dependence of cell cycle regulation on matrix stiffness in other cell types, we conducted similar experiments with C2C12 and 3T3 cells. From the fold change plots and BrdU results, we found that that unlike hMSCs, C2C12 cells showed a progressive increase in proliferation as we gradually increased the substrate rigidity. without any sharp transition (*Supplementary Fig. 4*). Additionally, they proliferated even when grown on very soft gels. For 3T3 cells, substrate stiffness had no effect on their proliferation (*Supplementary Fig. 5*).

### 3.(iv) Cell spread area does not correlate with cell cycle pattern vis-à-vis substrate stiffness but cellular traction does

Literature suggests that cell spread area and cellular traction increases with substrate stiffness, the former influencing DNA synthesis and reducing cell death [13][14][15]. To address which of these two factors correlate with observed substrate stiffness effect on cell proliferation, we analyzed them systematically. Results showed that the spread area of all three cell types increased gradually with substrate stiffness (Fig.2a-c). This pattern doesn’t match with the observed cell cycle behavior in which hMSCs displayed a drastic switch, C2C12 showed a gradual change, and 3T3 were insensitive to the substrate stiffness investigated.

Next, we addressed how contractile forces vary with varying substrate stiffness in the three cell lines. We found that hMSCs generated about 9.5 times higher contractile forces (around 320 ± 25 Pa) on a 5 kPa gel than on a 1 kPa gel (33 ± 3.37 Pa) (Fig. 2d). The C2C12 cells generated twice as much force on 5 kPa compared to 2 kPa. Note that on 1 kPa, C2C12 generated around 100 Pa traction force which is almost thrice as much as the forces generated by hMSCs on the same stiffness. Unlike the other two cell lines, traction force of 3T3 cells was much less sensitive for substrate stiffness. These trends mirror the fates of these cell types (proliferating and growth arrest) on substrates of various stiffness. Taken together, these observations suggest a possible involvement of cellular contractile proteins in sensing substrate stiffness and regulating cell cycle exit and re-entry in hMSCs.

**Figure 2.**
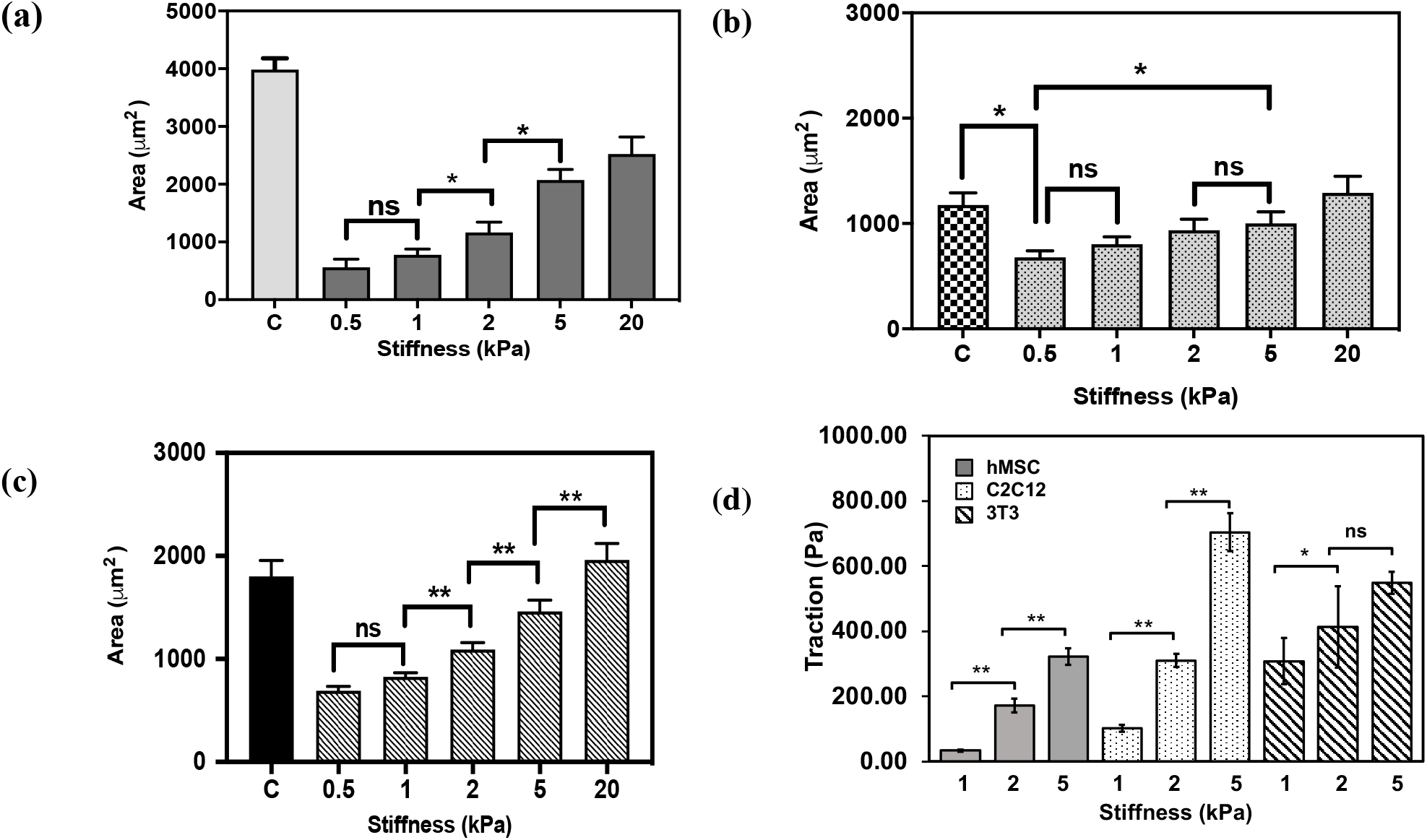
Effect of substrate stiffness on cell spread area and traction force in hMSC, C2C12 and 3T3 cells. Cell spread area across varying stiffness of PA gels and TCP (control) of hMSCs (a), C2C12 (b) and 3T3 cells (c) were calculated 24 hrs post-seeding. Result is represented as mean + SE, where n = 150 cells, N = at least 4 independent experiments, * p < 0.05, ** p < 0.005 and ns is non-significant (Student’s unpaired t-test). Also, traction versus stiffness for the three cell lines are plotted in (d). Result is represented as mean + SE, where n = atleast 10 cells per condition, N = at least 4 independent experiment.

### 3.(v) Substrate stiffness controls cell cycle of hMSCs via contractile proteins

To evaluate the possible involvement of contractile proteins in the regulation of cell cycle by matrix stiffness, we separately treated cells with ROCK inhibitor (Y-27632) and MEK-inhibitor PD98059. For the ROCK-inhibitor studies, a 5 kPa gel was chosen since the cells (a) proliferated here after their recovery from quiescence on softer substrates (Fig. 1), and (b) exerted significantly more traction force compared to softer substrates (Fig. 2d). The fold change and traction force values (*Supplementary Table 1*), obtained after treating with varying concentration of Y-27632, were normalized with their respective values for 5 kPa gel without the drug (Fig. 3). The same data for cells (untreated) grown on 1 kPa gel (Figs. 1 & 2), were plotted on the same graph (Fig. 3). Results show a gradual decrease of proliferation as well as cellular contractility with increased concentration of Y-27632 (Fig. 3a). For example, at 5 kPa with 20 μM Y-27632, cells were essentially quiescent and exhibited around 12% contractility compared to the control. On the other hand, at 10 μM, cells demonstrated 12% normalized contractile force and 15% normalized fold change, both measured against the control. These observations are similar to the effect of substrate stiffness on hMSCs behaviour. For example, cells grown on the softer 1 kPa gel become quiescent and exhibit 10% contractility with respect to the 5 kPa gel (Fig. 3a). Similarly, fold change assay carried out with PD98059 ERK inhibitor showed that cells on 5 kPa gel (treated condition) attained quiescent like condition similar to 1 kPa (Fig. 3b). As reported previously [16], our immunostaining results showed a substrate-stiffness specific response of p-ERK expression between 1 and 5 kPa gel, stiff substrate associated with significantly higher p-ERK expression (Fig. 3c & *Supplementary Fig. 6*). However, upon treatment with ERK inhibitor (∼1 μM) cells underwent a considerable reduction (∼21%) in p-ERK expression. ERK proteins are well known regulators of cell proliferation and contractility [17]. Our studies with ERK inhibitor also indicate possible involvement of ERK in substrate-stiffness induced regulation of cell proliferation. Taken together, our results with contractile inhibitors strongly imply that in hMSCs, substrate stiffness, cellular contractility, and proliferation are tightly coupled to each other and there is a positive correlation amongst them.

**Figure 3.**
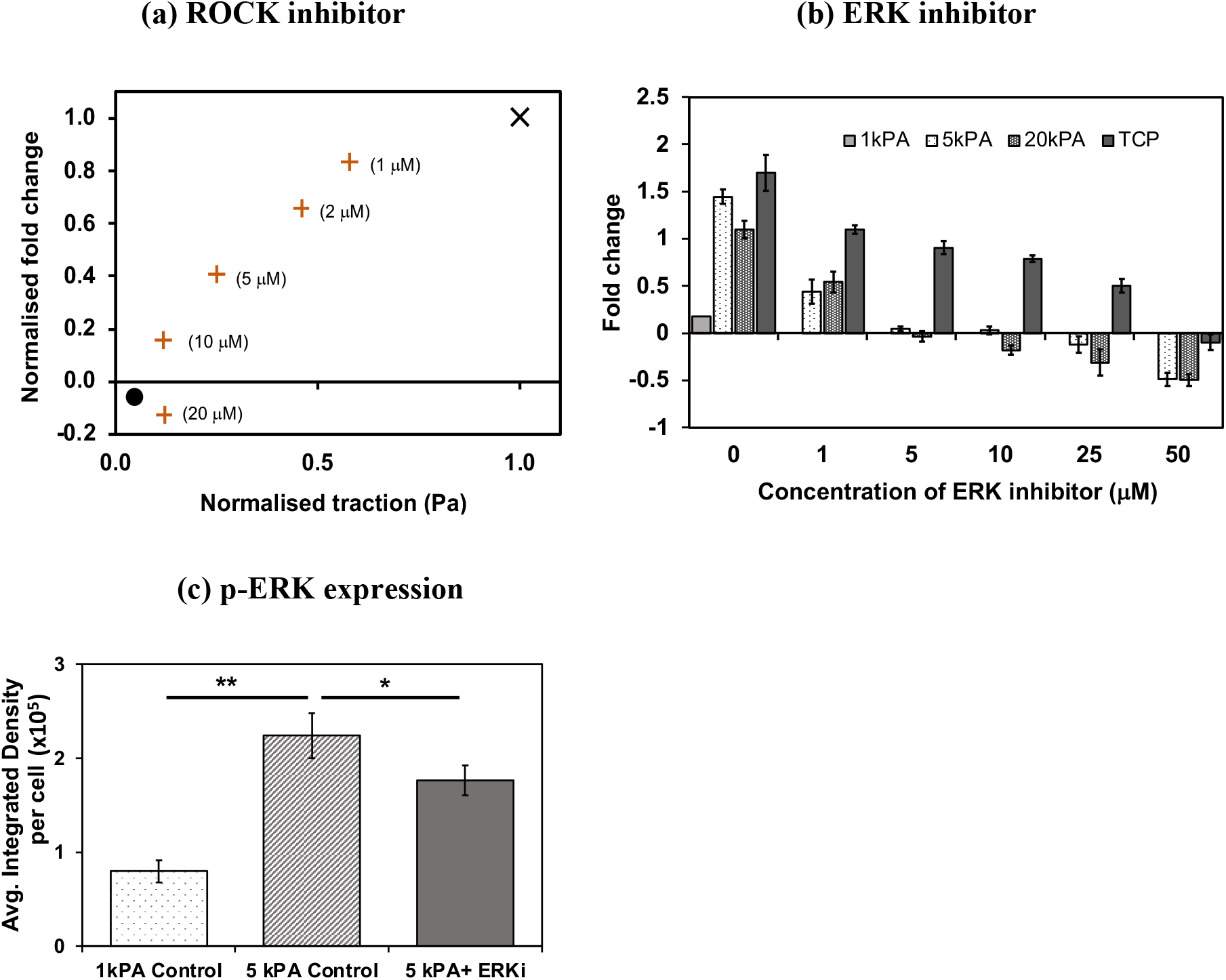
Cellular contractility regulates proliferation of hMSCs. (a) Cells are grown on 5 kPa gel and treated with varying concentration of the ROCK-inhibitor (Y-27632). Here, traction is on the abscissa, and ordinate gives the fold change proliferation for cells treated with Y-27632. ‘**+**’ represents values of cells grown on 5 kPa gel treated with varying doses of Y-27632, whereas ‘**×**’ represents cells on 5 kPa without the drug (control). Additionally, the fold change and traction force values for cells grown on 1 kPa gel, is indicated by ‘·’. All the values are normalized with their corresponding values for untreated (control) cells grown on 5 kPa substrate. Fold change values of cells treated with ERK-inhibitor are plotted in (b). Untreated cells on 1 kPa is used as a reference. Average integrated density of p-ERK expression in cells grown on 1 and 5 kPa gel (± ERK-inhibitor) are graphically shown in (c). ** p < 0.0001 and * p < 0.1 (Student’s unpaired t-test)

**Figure 4.**
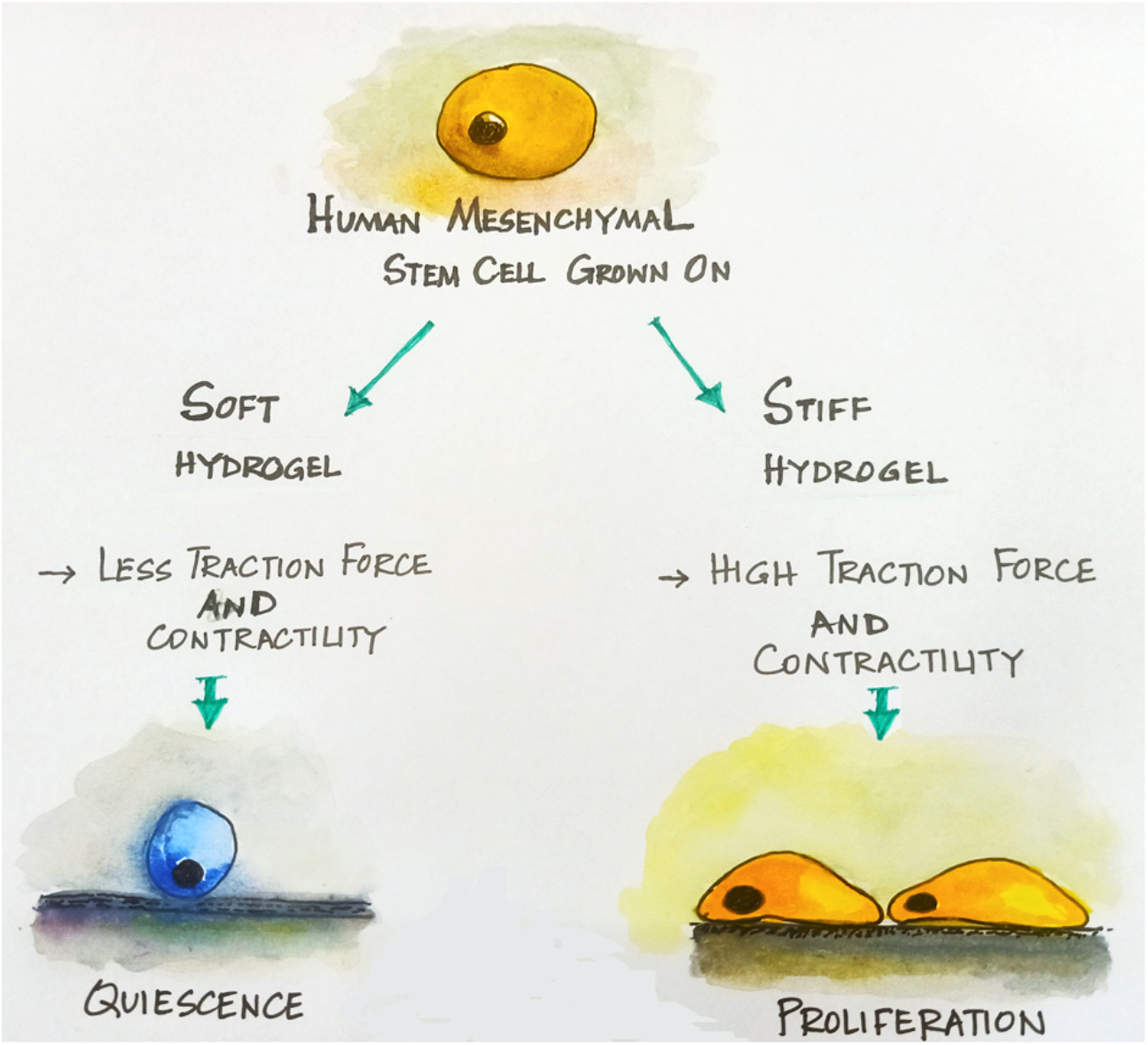
Graphical summary. Cellular contractility regulates proliferation of hMSCs. On soft substrate (≤ 1 kPa) human mesenchymal stem cells exert less contractile forces and remain in quiescent phase of cell cycle. On the other hand, on stiff substrate (≥ 5 kPa), they exert significant contractile forces and display prominent proliferation.

## 4. DISCUSSION

Winer et al. (2008) showed that stiffness of bone marrow (∼ 0.25 kPa) contributes to the maintenance of quiescent multipotent hMSCs [4] which were quiescent state on soft gel (0.25 kPa) but proliferating on stiff substrate (7.5 kPa PA gels). This study established the unique role of ECM stiffness on the maintenance of quiescent and proliferating pool of multipotent stem cells *in-vitro*. However, neither did it provide the precise range of substrate stiffness that switches the cell cycle of stem cells “on” or “off”, nor did it analyse the regulatory factors that govern them. It is important to address these lacunae to have a precise understanding of how changes in ECM is associated with cell behaviour. In this work, we evaluated a range of substrates stiffness – viz. 0.5, 1, 2, 5, and 20 kPa – and report a noticeably sharp transition from quiescence to proliferation between 1 and 5 kPa for bone-marrow derived hMSCs. To understand whether the above nature of cell response towards substrate stiffness follows a universal pattern, we examined C2C12 and 3T3 cell lines across the same range of stiffness. Interestingly, we found differences in proliferation profiles amongst all the cell lines. Further examination of cell spread area and cellular contractility revealed that contrary to earlier findings [13], the cellular contractility, but not the cell spread area, were corelated with the sensitivity of the proliferation of hMSCs, C2C12, and 3T3 cells towards substrate stiffness.

In an earlier study, it has been shown that the fates of C2C12 cells are largely determined by the mechanical cues of the environment[18]. Cells in a 2-D culture, proliferated independent of the stiffness of alginate gels compared to the primary myoblasts that displayed substrate-stiffness sensitive proliferation, whereas in a 3-D culture C2C12 cells displayed stiffness sensitivity. Contrary to these observation on a 2-D culture, we find in our study that the C2C12 cells display a stiffness-dependent growth without attaining quiescence on the softest gel (0.5 kPa) chosen for our study. The lack of correlation between the two observations may be due to two different gel systems used. On the other hand, the 3T3 fibroblasts remained non-responsive to the changes in this range of matrix stiffness.

On stiff substrates, cells typically display prominent stress fibres, exhibit a more spread phenotype, and modify the properties of adhesion and contractile proteins. These changes eventually increase the stress applied by cells on the underlying substrates, and subsequently regulate the activity of small GTPases and formation of focal adhesions[2]. Well spread cells on stiff substrates, when treated with inhibitors of cellular contractility undergo a disassembly of stress fibres and reduction in cell proliferation. Restricting cell spread area alone also reduces cell proliferation[19]. Therefore, both cell spread area and cellular contractility are tightly coupled to cell proliferation. From another study, it is also known that cell spread area and substrate stiffness are significant predictors of cellular contractility[20]. Altogether these findings suggest that cell proliferation, cellular contractility, cell spread area and matrix stiffness are functionally linked. To determine how substrate stiffness is coupled to cell proliferation in our study, we investigated changes in cell spread area and cellular contractility across substrates of varying stiffness. Our results demonstrated that all the three cell lines responded to changes in substrate stiffness in varying degrees in terms of their spread area. However, the trends captured did not correlate with their corresponding proliferation profiles. This leads to the possible involvement of other factors, like cellular contractile proteins, in regulation of cell proliferation. Interestingly we observed that the cellular contractility trends in these cell lines corresponded to their growth curves (fold change values). In particular, hMSCs displayed a sharp increase in their contractile forces between 1 & 5 kPa that correlated to their characteristic proliferation profile, i.e., quiescence (1 kPa) & growth (5 kPa). Cell cycle duration, and phase transition events are largely governed by the cellular forces generated by them[21]. It is observed that the tensile forces generated by cells vary in a bi-phasic manner during cell cycle; i.e. they increase (G1), reach a plateau (S-phase) followed by a decline (G2 phase)[22]. Reduction in cellular tension is associated with cell cycle exit. Our results with hMSCs reflect this pattern, displaying bi-phasic curves of both contractility and proliferation.

Since amongst the three cell lines, hMSCs responded to substrate stiffness in a biphasic manner, we conducted further studies using inhibitors of ROCK and MEK-signalling pathways, both having known involvements in regulating cell proliferation, adhesion, migration, etc.[23][24][25][26]. Interestingly we found a dose-dependent inhibitory effect of both the inhibitors of cellular contractility on hMSC proliferation. Cells on 5 kPa gel and treated with varying doses of ROCK inhibitor (Y-27632) underwent gradual reduction in both their fold change values and contractile forces. Between 10 and 20 μM of Y-27632 cells attained 1 kPa like-quiescent condition. It has been reported in literature that p-ERK displays a substrate-stiffness sensitive expression[16]. However, the expression level of p-ERK between 1 & 5 kPa PA gel was not evaluated. Our results showed that although the expression of p-ERK on 5 kPa gel is approximately 2.5 folds higher than 1 kPa gel, with ERK inhibitor, the expression was suppressed (∼ 21%) which subsequently reduced the fold change of proliferation of cells.

Our study therefore suggests the involvement of contractile proteins in translating extracellular matrix stiffness changes to cell interior and regulation of cell cycle of hMSCs. One possible mechanism is – Rho and/ROCK proteins respond to the mechanical cues and via mediators of cellular contractility like FAK, activate Cyclins D1 and A, eventually regulate different phases of cell cycle[27].

## 5. CONCLUSION

Our study demonstrates that substrate stiffness regulates the proliferating and quiescence phases of cell cycle in hMSCs via cellular contractility. Further studies addressing the mechanism(s) associated with this process will offer new mechanistic insights into stem cell research for therapeutic advances.

## AUTHOR CONTRIBUTIONS

A.M., S.K.K, & S.S. designed the experiments. S.K.K, S.S, P.P, & S.B. performed the experiments and analyzed the data. A.M. and S.S. prepared the manuscript.

## Acknowledgement

AM acknowledges the Wadhwani Research Centre for Bioengineering, IIT Bombay, India and Wellcome Trust-DBT India Alliance (Project #IA/E/11/1/500419) for research support. SS thanks IIT Bombay for her Post-Doctoral Fellowship. We thank Dr. James P Butler (Harvard Medical School, Department of Medicine, Boston) for his TFM codes, and Dr. Jyotsna Dhawan, InStem, for generously donating the C2C12 and 3T3 cells.

## Supplementary Figures and table

**Supplementary Fig.1:**
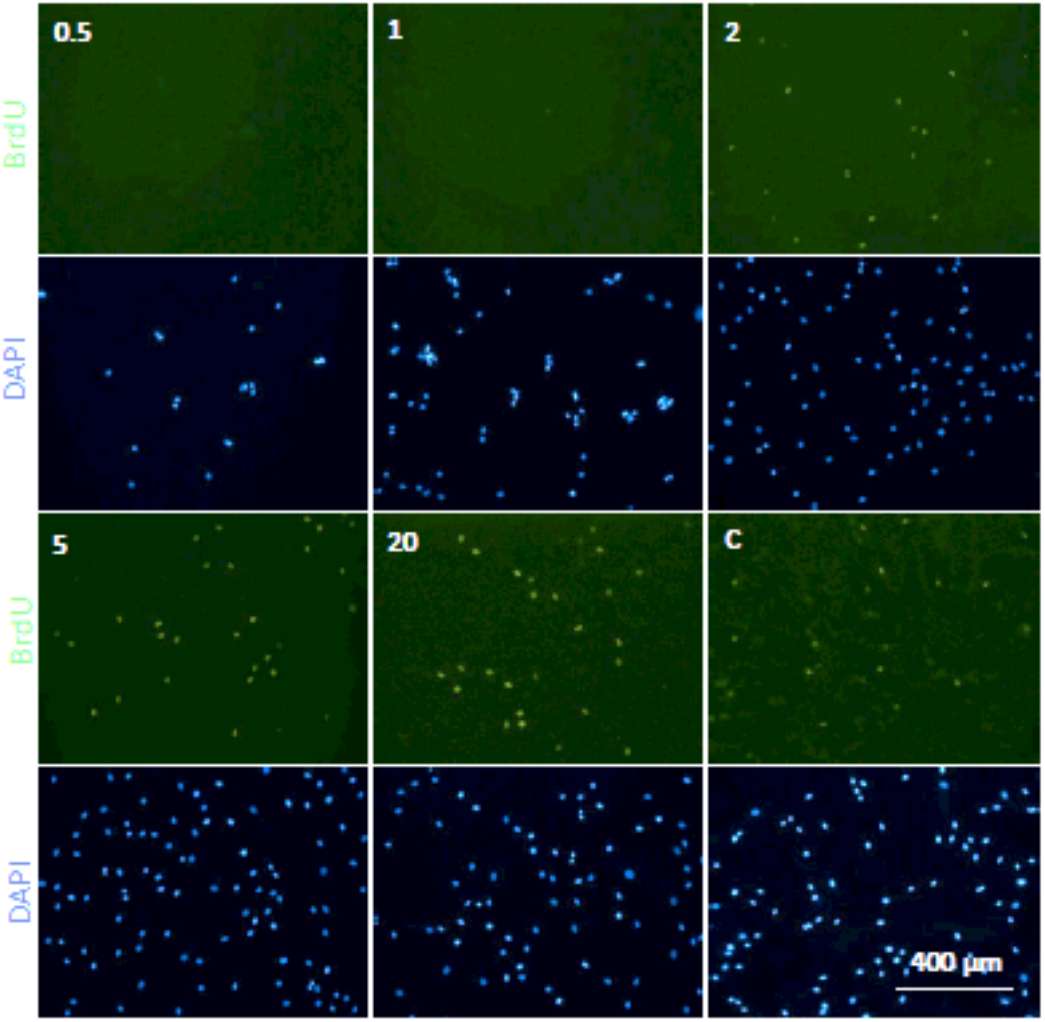
Effect of substrate stiffness on S-phase of hMSC using BrdU assay. (a) Representative images of BrdU positive cells at varying stiffness (Green: BrdU, Blue: DAPI). Scale bar is 400 μm (b) Plot of percentage of BrdU positive cells vs stiffness and control (C). Result is represented as mean + SE, where n = 300 cells, N = 3 independent experiments, * p < 0.05 (Student’s unpaired t-test) and ns is non-significant.

**Supplementary Fig.2:**
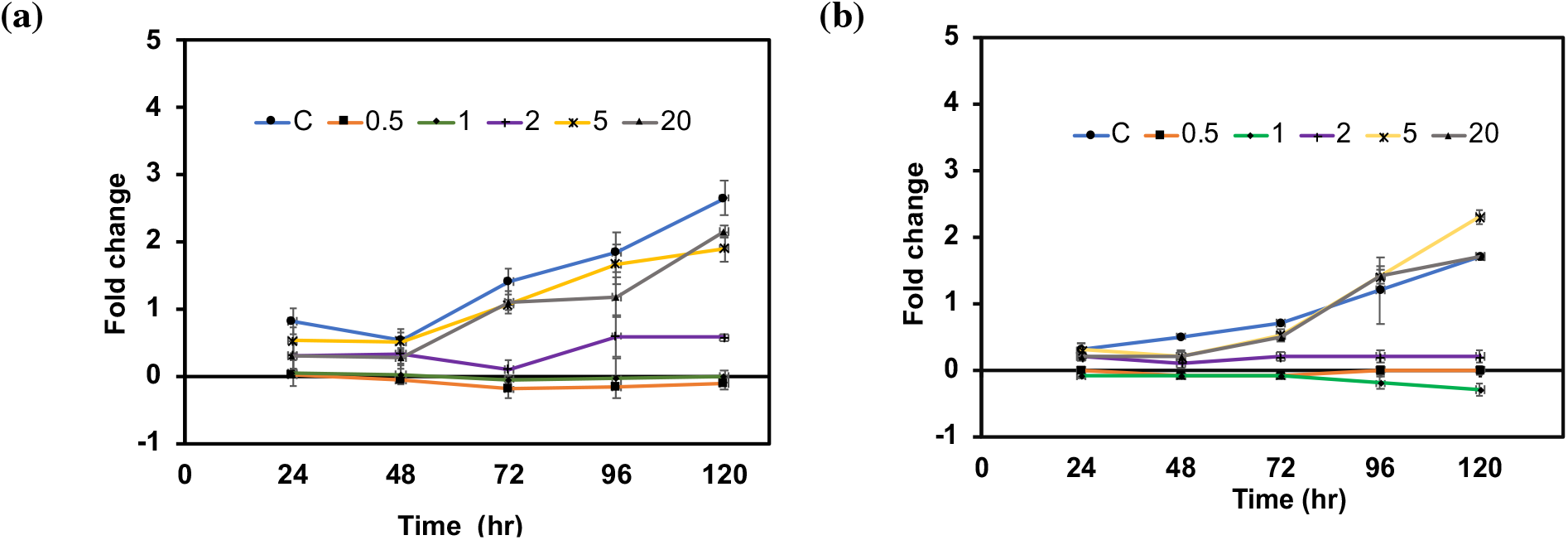
Effect of substrate stiffness on exit and re-entry of hMSCs. (a) Fold change vs time plot for cell cycle exit and re-entry of hMSCs (b). In both the graphs experimental conditions are compared with the TCP control (C). Result is represented as mean + SE, where N = 3 independent experiments.

**Supplementary Fig.3:**
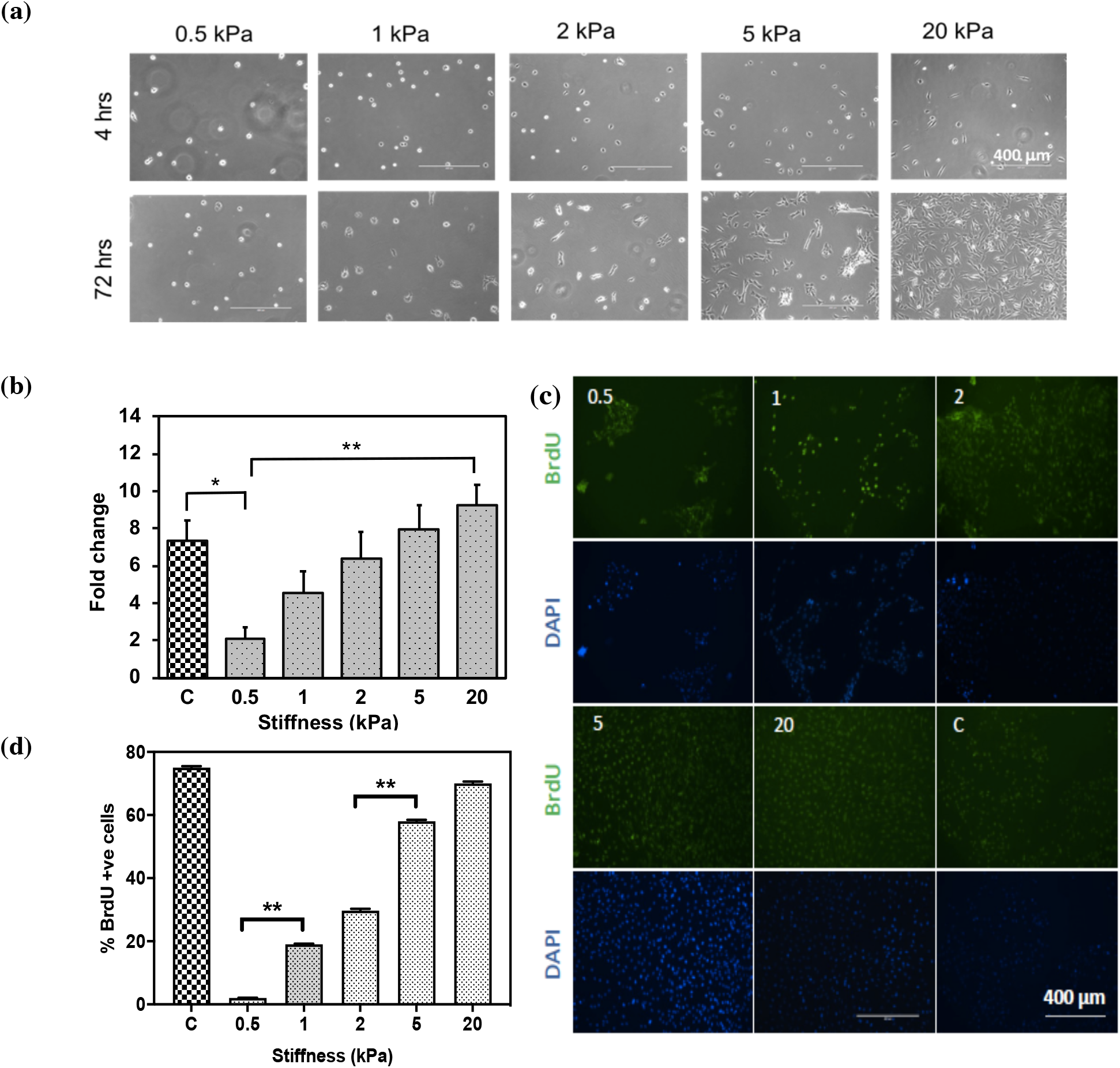
Effect of substrate stiffness on cell cycle exit of C2C12 by fold change and BrdU assays. (a) Representative images of cells on PA gels of various stiffness captured after 72 hrs. Scale bar is 400 μm (b) Fold change vs stiffness plot at 72 hrs. Result is represented as mean + SE, where N = at least 4 independent experiments, * p < 0.05 (Student’s unpaired t-test). (c) Representative images of BrdU positive cells at varying stiffness and TCP. (Green: BrdU positive, Blue: DAPI) (d) Plot for BrdU positive cells vs stiffness. In both the graphs, experimental conditions are compared with the TCP control (C). Result is represented as mean + SE, where n = 300, and N = at least 4 independent experiments, ** p < 0.005 (Student’s unpaired t-test).

**Supplementary Fig.4:**
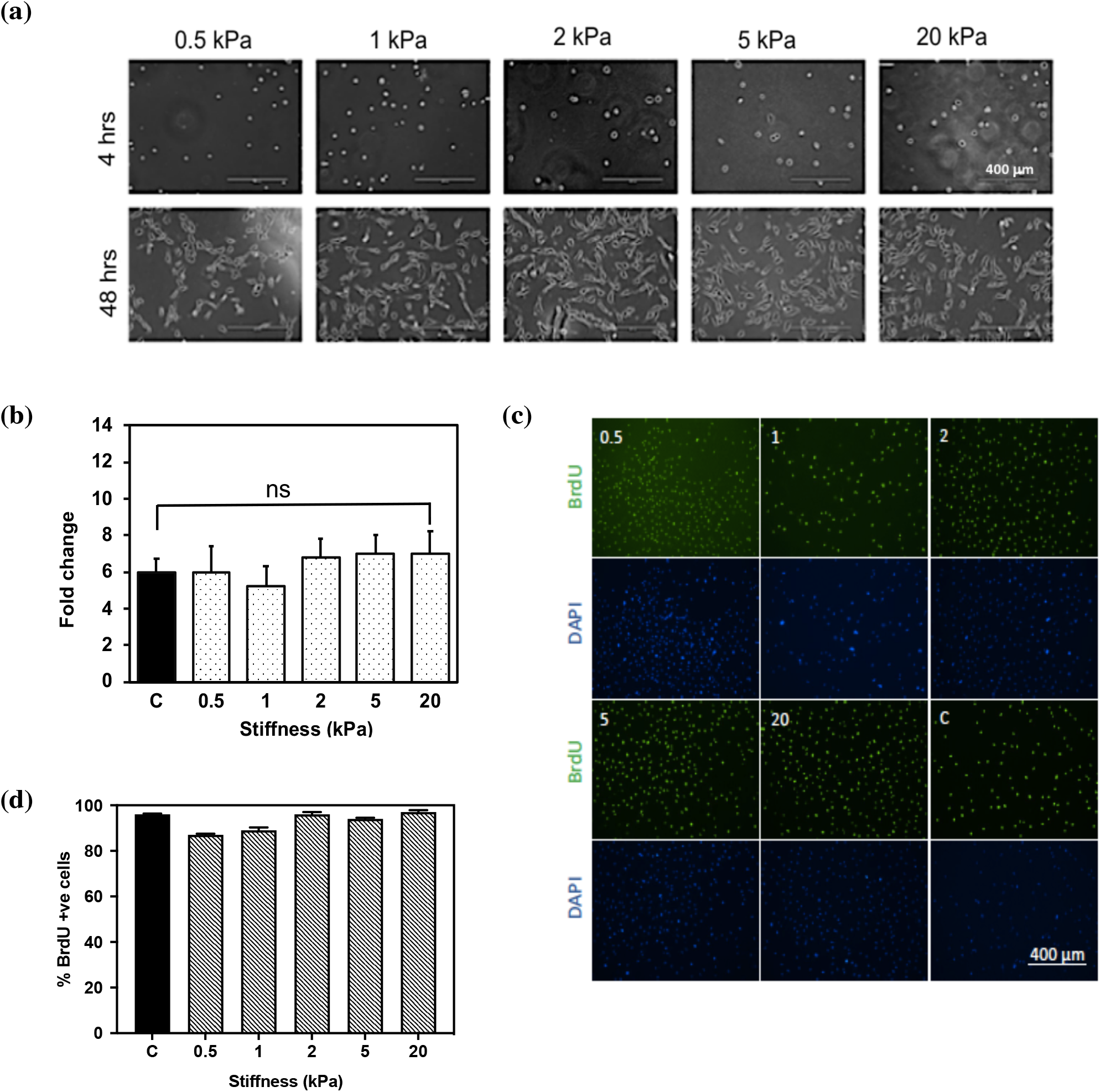
Effect of substrate stiffness on cell cycle exit of 3T3 cells by fold change and BrdU assays. (a) Representative images of 3T3 on substrate of varying stiffness after 72 hrs respectively. Scale bar is 400 μm. (b) Fold change versus stiffness plot for 3T3 at 72 hrs. Result is represented as mean + SE, where N = 4 independent experiments (c) Representative images of BrdU positive cells at varying stiffness and TCP control. Green and blue dots indicate BrdU positive and DAPI stained nuclei (d) Plot for BrdU positive cells vs stiffness. In both the graphs experimental conditions are compared with the TCP control (C). Result is represented as mean + SE, where n = 300, and N = at least 4 independent experiments, ns is non-significant.

**Supplementary Fig.5:**
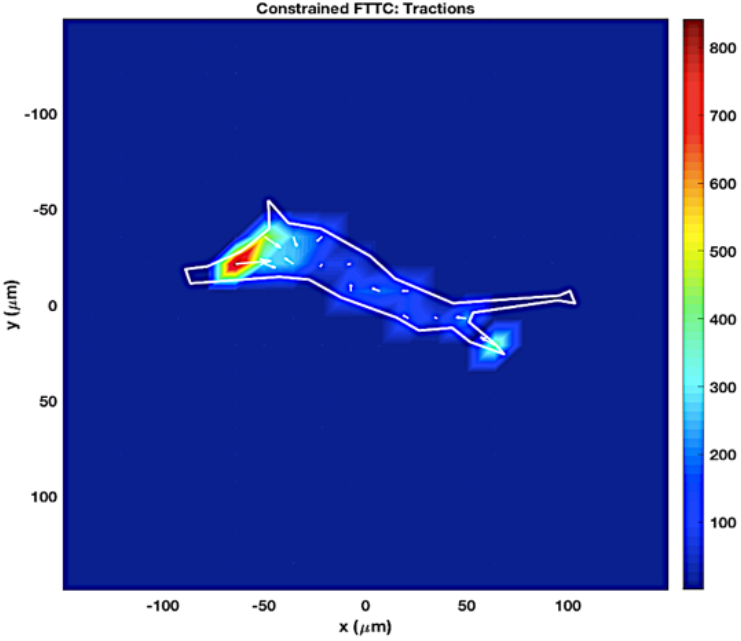
Representative traction stress map obtained from traction force microscopy (TFM) of hMSC on a 5 kPa gel during stressed condition.

**Supplementary Fig.6:**
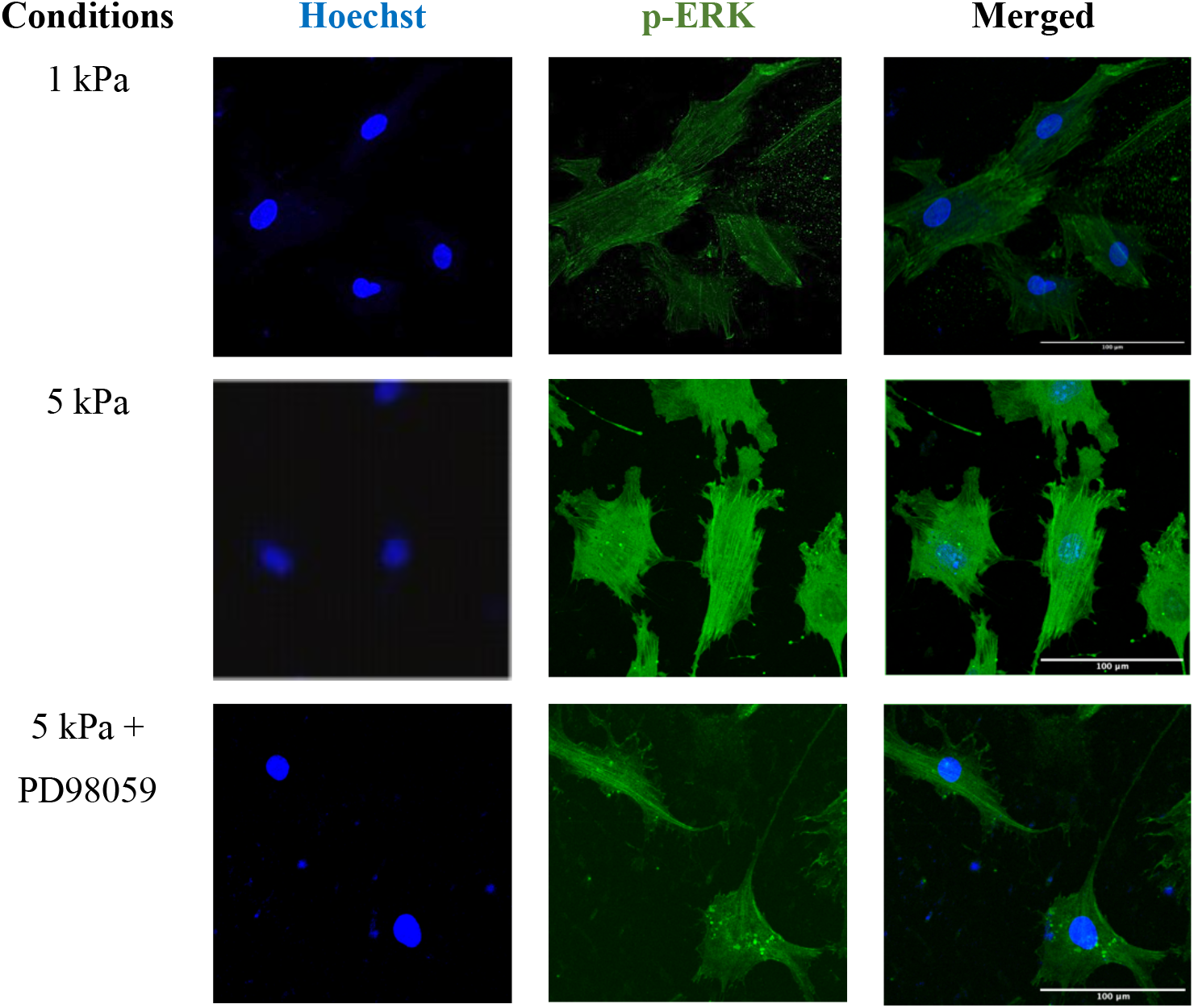
Immunostaining of hMSCs with p-ERK antibody. Representative images of cells grown on 1 and 5 kPa (with and without ERK-inhibitor treatment for 48 hrs).

**Table 1:** Values of contractile forces (Pa) generated by cells with the treatment of ROCK-inhibitor (Y-27632) of varying doses on 5 kPa gel

| Conditions (± Y-27632) on 5 kPa PA gel | Traction force (Pa) |
| --- | --- |
| Control | $322 \pm 24$ |
| 1μM | $186 \pm 11$ |
| 2μM | $148 \pm 28$ |
| 5μM | $80 \pm 9$ |
| 10μM | $38 \pm 5$ |
| 20μM | $39 \pm 5$ |

Results are represented as avg. traction force ± SEM from at least 4 independent experiments and analysing at least 26 cells per condition.

